# Latent transcriptional programs reveal histology-encoded tumor features spanning tissue origins

**DOI:** 10.1101/2023.03.22.533810

**Authors:** Hanna M. Hieromnimon, James Dolezal, Kristina Doytcheva, Frederick M. Howard, Sara Kochanny, Zhenyu Zhang, Robert L. Grossman, Kevin Tanager, Cindy Wang, Jakob Nikolas Kather, Evgeny Izumchenko, Nicole A Cipriani, Elana J. Fertig, Alexander T Pearson, Samantha J Riesenfeld

**Affiliations:** Graduate Program in Biophysical Sciences, University of Chicago, Chicago, IL; Department of Medicine, University of Chicago, Chicago, IL; Department of Pathology, University of Chicago, Chicago, IL; Center for Translational Data Science, University of Chicago, Chicago, IL; Else Kroener Fresenius Center for Digital Health, University Hospital Carl Gustav Carus Dresden, Technical University Dresden, Dresden, Germany; Department of Medical Oncology, National Center for Tumor Diseases, University Hospital Heidelberg, Heidelberg, German; Division of Pathology and Data Analytics, Leeds Institute of Medical Research at St. James’s, University of Leeds, Leeds, United Kingdom; Departments of Oncology, Biomedical Engineering, and Applied Mathematics and Statistics, Johns Hopkins University, Baltimore, MD; Convergence Institute, Johns Hopkins University, Baltimore, MD USA; Pritzker School of Molecular Engineering, University of Chicago, Chicago, IL; Committee on Immunology, University of Chicago, Chicago, IL; Institute for Biophysical Dynamics, University of Chicago, Chicago, IL

## Abstract

Precision medicine in cancer treatment depends on deciphering tumor phenotypes to reveal the underlying biological processes. Molecular profiles, including transcriptomics, provide an information-rich tumor view, but their high-dimensional features and assay costs can be prohibitive for clinical translation at scale. Recent studies have suggested jointly leveraging histology and genomics as a strategy for developing practical clinical biomarkers. Here, we use machine learning techniques to identify *de novo* latent transcriptional processes in squamous cell carcinomas (SCCs) and to accurately predict their activity levels directly from tumor histology images. In contrast to analyses focusing on pre-specified, individual genes or sample groups, our latent space analysis reveals sets of genes associated with both histologically detectable features and clinically relevant processes, including immune response, collagen remodeling, and fibrosis. The results demonstrate an approach for discovering clinically interpretable histological features that indicate complex, potentially treatment-informing biological processes.

## Introduction

Squamous cell carcinomas (SCCs) represent a diverse subtype of cancers. Squamous cells form a layer within the skin and mucus membranes of organs, including the lung, head and neck, and cervix. The mechanisms that cause these cells to transform into cancer overlap across different primary organ systems (“primary”). For example, the oncogenic human papillomavirus (HPV) occurs in cervical (CESC)^1^ and head and neck (HNSC) SCC^2^, while smoking represents a common etiology of lung SCC (LUSC)^3^ and HNSC^2^. Additionally, previous analyses of transcriptional profiles in The Cancer Genome Atlas (TCGA) demonstrate substantial similarity across LUSC, HNSC, and CESC datasets^4^. Given their overlapping etiologies, transcriptional similarities, and common cell type, SCCs are ideal for analyzing common and tissue-specific processes in a large set of cancers.

Precision medicine approaches to cancer treatment rely on understanding the variables driving key biological processes in tumorigenesis, such as proliferation, invasion, and immune evasion. Identifying biomarkers for these processes is requisite for improving therapeutic strategies and identifying patients who will benefit from targeted therapies. A major barrier to understanding these variables has been their complexity and dependence on the genomic, metabolic, and other microenvironmental influences over the tumor^5^. Prior work has shown that cancer type and subtype, as well as established disease-altering variables, such as HPV status, are encoded in latent space embeddings of RNA-seq^6,7^, methylation^8^, and multi-omic^9,10^ bulk profiles of cancer tissue. Single-cell molecular profiling and immune-subtype profiling have also identified recurrent phenotypes^11–13^ across cancers. However, a data-driven, multi-modal analog of Hanahan and Weinberg’s conceptual “hallmarks of cancer”^14^ has yet to be fully established and related to treatment.

A further challenge in clinical deployment of gene-expression based biomarkers^12,15–17^ is the time, expertise, and monetary cost associated with collecting and analyzing this data type^18^. In contrast, H&E slides are routinely collected as a part of the standard of care in cancer diagnosis and are ubiquitously available. Tumor histology can reveal additional prognostic information, including genomic features. Indeed, recent work used deep learning models to predict genomic alterations and gene expression data from tumor histology^19–21^; other efforts combined genomics and histology to predict prognosis^22^, overall survival^23^, and, leveraging new spatial transcriptomics approaches, molecular heterogeneity of cell states^24^. While genomics and histology offer complementary, synergistic insights into latent tumor biology, clinical deployment of deep-learning-based histology biomarkers requires model transparency demonstrating the histologic features responsible for predictions^25^.

Here, we present a gene-expression derived CoGAPS^26^ latent factor (CLF) analysis of SCCs, highlighting key known and novel features of the disease. This unsupervised, comprehensive analysis unites established genomic, clinical, and histologic features of SCCs. We used deep learning to both detect CLFs in tumor histology and generate synthetic histology images, subsequently annotated by a pathologist, with histologic features associated with the CLFs. To train our model, we compiled RNA-seq and histological data from 1,300 lung, head and neck, and cervical SCCs from TCGA, and then validated the latent space using RNA-seq data from 174 lung and head and neck SCCs in the Clinical Proteomic Tumor Analysis Consortium (CPTAC) database. Our results provide insight into the biological variables that vary within SCCs and provide a methodology for translating this insight into the clinical setting using readily available H&E slides.

## Results

### Identification of transcriptional CoGAPS latent factors (CLFs)

To identify gene-expression derived CLFs, we performed latent space factorization on log-scaled TMM-normalized gene expression data from HNSC, CESC, and LUSC TCGA samples (n = 1300) (**Fig. 1a**). CoGAPS^26^ decomposed the initial gene expression matrix, indexed by genes (rows) and samples (columns), into the product of two lower-dimensional matrices: the pattern matrix, indexed by genes (rows) and CLFs (columns); and the amplitude matrix, indexed by CLFs (columns) and samples (rows). The number (12) of CLFs was selected as a reasonable tradeoff between higher values, which captured more programs, both within and across the three primary tissues (**Table 1**), and lower values, which gave better coherence (e.g., in amplitudes on a UMAP) and more unique top-weighted genes in the pattern matrix (**Supp. Fig. 1**).

**Table 1:** CoGAPS latent factors (CLFs) from transcriptomics matrix factorization identify biological programs with multiple levels of supporting evidence. Table of CLFs with suggested labels, supporting associated genes, enriched biological pathways, clinical and gene signature associations, and pathologist-characterized histologic features.

|  | Suggested Label | Genes | Pathways | Clinical | Histology |
| --- | --- | --- | --- | --- | --- |
| Factor 1 | DNA repair/Cervical | SYCP2, CDKN2A, CDCA7 | DNA Repair, HRD Through Homologous Recombination HRR | Cervical | Smaller nuclei, darker cytoplasm, diminished stroma, increased lymphocytes |
| Factor 2 | HPV positive |  |  | HPV positive | Hyperchromatic nuclei, diminished stroma |
| Factor 3 | Epithelial cells | KRT7, WFDC2, SCNN1A | Defective GALNT3 causes HFTC, Dectin-2 family |  |  |
| Factor 4 | Cellular stress | RPL13, RPL8, RPLP1 | Cellular response to starvation |  |  |
| Factor 5 | B cell immune | IGKC, IGHA1, IGHG1, IGHG3, IGHG2, JCHAIN | Signaling by the B Cell Receptor (BCR), Initial Triggering of Complement |  | Increased lymphocytes |
| Factor 6 | T cell immune | HLA-DRA, HLA-DRB1, HLA-DPA1, CD74, HLA-DQA1, CXCL9, CXCL10 | Cytokine Signaling in Immune System, PD-1 signaling | IFNG/IMS gene signature | Increased stroma, increased lymphocytes |
| Factor 7 | Lung |  |  | Lung |  |
| Factor 8 | Fibrosis | COL1A2, FN1 | Extracellular matrix organization, Collagen degradation | Cancer Associated Fibroblast gene signature |  |
| Factor 9 | Smoking status |  | Biological oxidations | Smoking history | Increased, prominent nucleoli or replaced by necrotic material/lymphocytes |
| Factor 10 | Metabolism | GAPDH, HSPA8, HSP90AB1 |  |  |  |
| Factor 11 | Collagen remodeling | KRT17, KRT6A, KRT5, MMP1, LAMC2, LAMB3, LAMA3 | Collagen formation, Programmed Cell Death | pEMT gene signature | New tumor cells appear focally, focally tumor cells decrease in number, nucleoli become larger, chromatin is open, prominent nucleoli, increased cytoplasm, increased stroma, decreased lymphocytes |
| Factor 12 | Head and Neck |  | Keratinization, Formation of the Cornified Envelope | Head and Neck |  |

**Figure 1:**
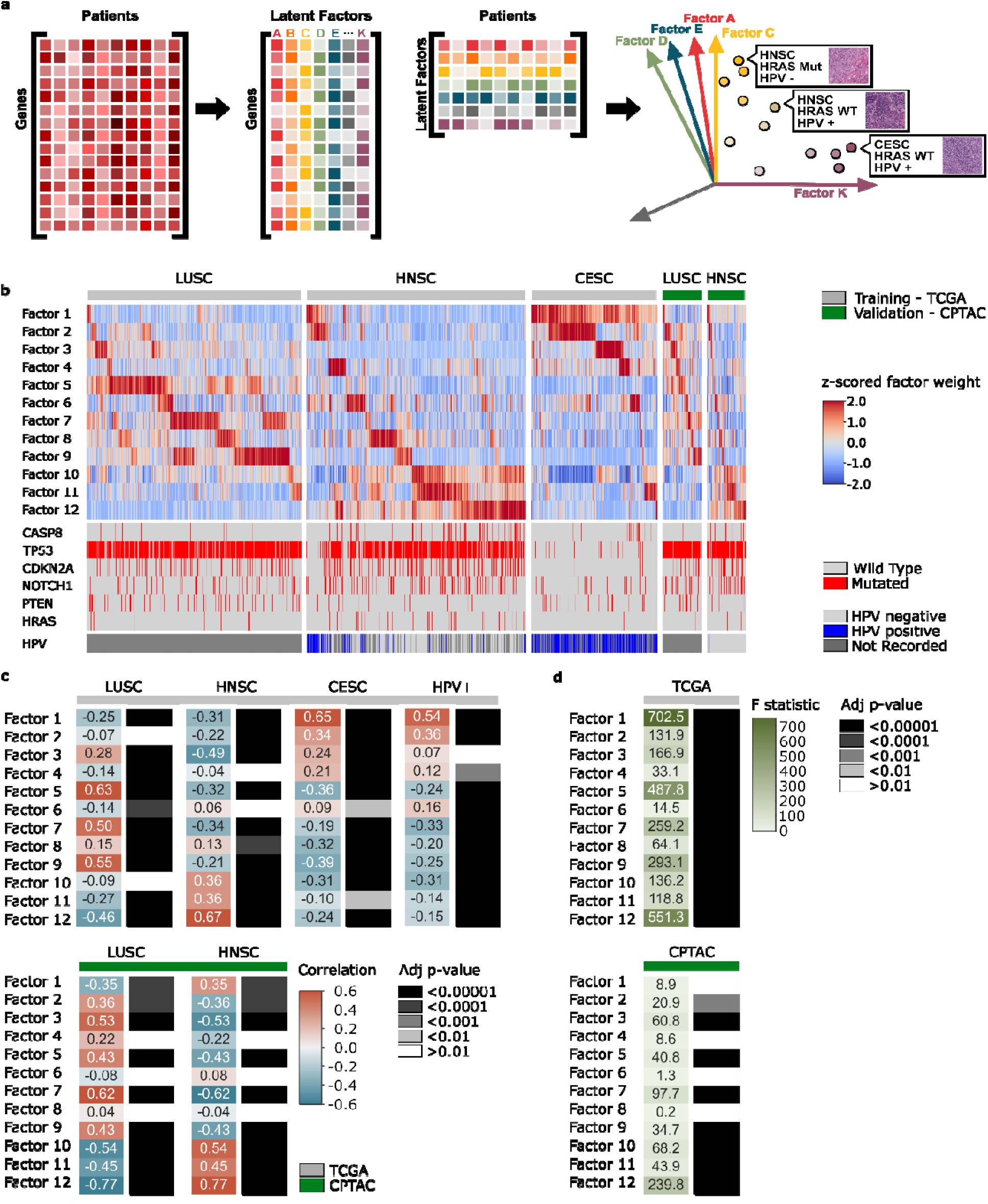
Transcriptomic latent space factorization characterizes inter- and intra-primary tumor variation. **a**, Schema of the CoGAPS factorization of the transcriptomics matrix (left) into two lower-dimensional matrices (middle), parameterized by the number of latent factors; each matrix is then analyzed in the context of other ‘omics and histology image data (right). **b**, Heatmap shows sample weights (color) of CoGAPS latent factors (upper rows), reported mutation states (color) of cancer-associated genes (middle rows), and infection status (color) for human papilloma virus (HPV, bottom row) in patients (columns) with primary squamous cell (SC) carcinomas (headings; LUSC: lung; HNSC: head and neck; CESC: cervical), from two datasets (top row annotation; TCGA: the Cancer Genome Atlas; CPTAC: the Clinical Proteomic Tumor Analysis Consortium). CPTAC data are projected into TCGA-inferred latent space. Weights z-scored by row and dataset. **c**, Spearman’s rho correlations (left, color) between weights for latent factors (rows) and primary or HPV-status groups (headings), with Bonferroni-corrected p-values (right, gray scale), in TCGA (top) and CPTAC (bottom). **d**, Variability (ANOVA F statistic, left, color) in weights of latent factors (rows) across primary types; darker color indicates more tissue-specific variation; with Bonferroni-corrected p-values (right, gray scale).

CLF weights created groupings of patients compatible with major clinical variables and known differentiators of disease (**Supp. Fig. 2**), demonstrating that CLFs capture key aspects of the underlying biology. As expected, some factors were enriched in specific primary tissues, i.e., CLF 1 in cervical (“*Factor 1 – Cervical”*), CLF 7 in lung (“*Factor 7 – Lung”*), and CLF 12 in head and neck (“*Factor 12 – Head and Neck”*) cancers (**Fig. 1b-d**). We validated their tissue specifity by projecting the log-scaled TMM-normalized gene expression data from SCCs available in CPTAC (LUSC and HNSC) into the space of TCGA-derived CLFs (**Fig. 1b**). Primary-specific CLFs “*Factor 7 – Lung”* and “*Factor 12 – Head and Neck”*, identified from TCGA, aligned with the corresponding primaries in the independent CPTAC dataset. Additionally, in z-score-normalized CLF embeddings of TCGA and CPTAC data, samples of the same primary type tend to be neighbors (**Supp. Fig. 3**). Beyond primary-specific CLFs, the remaining CLFs encoded across-primary programs spanning tissue origins (**Fig. 1b–d, Table 1**).

### Transcriptional CLFs align with salient biological programs

High gene weights (pattern weights) for a specific CLF indicate a gene expression program that tends to be highly expressed in patients with high sample (amplitude) weights for that CLF. To study the gene expression programs associated with CLFs, we investigated the top-weighed genes in each CLF (**Supp. Fig. 4**). For example, CLF 1 has high weights for DNA replication and repair genes *SYCP2, CDKN2A*, and *CDCA7*, leading us to refine its label accordingly (*“Factor 1 – DNA Repair/Cervical”*). Epithelium-associated genes (*KRT7, WFDC2, SCNN1A, KRT8*) are highly weighted in CLF 3 (“*Factor 3 – Epithelial cells”*), while genes encoding ribosomal proteins, previously shown to differentiate between TCGA tumors^27^, (*RPL13, RPL8, RPLP1*) suggest a cellular stress program in CLF 4 (“*Factor 4 – Cellular stress”*). CLF 6 is comprised of major histocompatibility complex genes (*HLA-DRA, HLA-DRB1, HLA-DPA1, CD74, HLA-DQA1*) and chemokines (*CXCL9, CXCL10*), indicating a gene expression program related to antigen presentation and T cell infiltration (“*Factor 6 – T cell immune”)*. In contrast to that immune program, CLF 5 is characterized by a humoral and B-cell mediated response, with high expression of immunoglobulin genes (*IGKC, IGHA1, IGHG1, IGHG3, IGHG2*) and joining chain of multimeric IgA and IgM (*JCHAIN*) (“*Factor 5 – B cell immune”*). High weights for fibroblast markers *COL1A2* and *FN1*, according to the Genotype-Tissue Expression (GTEx) project (**Supp. Fig. 5**), suggest CLF 8 may capture more fibrotic tumors (“*Factor 8 – Fibrosis”*). Oxidative stress (*GPX2*) and tumor initiation (*SOX2*^28^, *TP63*^*29*^) are highly weighted in CLF 9, suggesting a potential smoking etiology related factor^30^ (“*Factor 9* – *Smoking status”*). CLF 10 has high weights for metabolism and stress-response genes (GAPDH, HSPA8, HSP90AB1) (“*Factor 10 – Metabolism”*). Finally, keratin genes (*KRT17, KRT6A, KRT5*), matrix metallopeptidase genes (*MMP1*), and laminin genes (*LAMC2, LAMB3, LAMA3*) are highly weighted in CLF 11 (“*Factor 11 – Collagen Remodeling”)*, suggesting a gene expression program related to extracellular matrix modification and localized tumor invasion. Together with the described enrichment of *Factors 1, 7*, and *12* in specific primaries, these data support a hypothesis of biological functions associated with each CLF.

To further interrogate the biological programs captured by CLFs, we performed pathway enrichment analysis using g:Profiler (g:GOSt)^31^ on the top 100 weighted genes in each CLF. From the set of significantly enriched pathways (FDR-adjusted *P* < 0.01) for a CLF, we manually selected a succinct list of biologically distinct pathways (**Supp. Table 1**). To better interpret CLF-associated functions, we focused on pathways in this set whose gene expression was significantly associated with sample weights for the CLF. Specifically, we identified those pathways for which the average expression of all pathway genes was significantly different between sets of patients classified as “low” or “high” for the CLF, depending on whether the sample weight was below the 60th or above the 75th percentile, respectively, of weights for that CLF(**Fig. 2a**).

**Figure 2:**
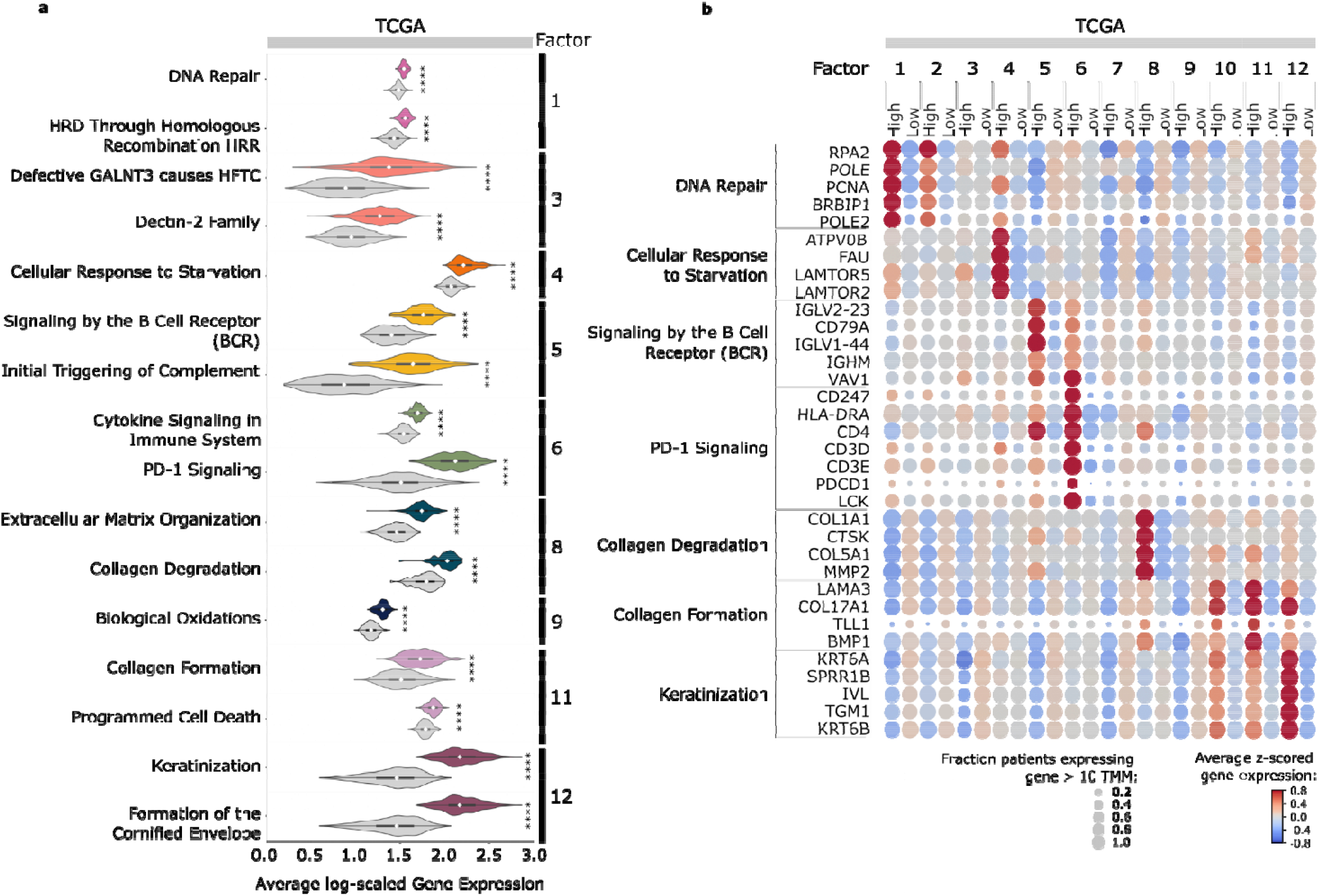
Latent factor genes are enriched in distinct, cancer-associated pathways. **a**, Violin plots show the distributions of the average log expression of all genes in a pathway (row label) that was identified as enriched in genes associated with the indicated latent factor (label on right), across TCGA patients who have high (in color) or low (in gray) weights for that factor. Internal boxplot shows the median (white diamond), the first quartile and third quartile (left and right ends of thicker box), and the minimum and maximum (left and right ends of thin line). Distributions compared by *t*-test (**** *P* < 0.0001) **b**, Dot plot shows the average expression (color) of individual genes z-scored by expression across the dataset in TCGA patients, grouped by high or low factor weight (columns, annotations above), for select genes (rows) in enriched pathways (annotations on left). Dot size indicates the fraction of patients in the group expressing the gene.

The analysis revealed that “*Factor 1 – DNA Repair/Cervical”* “high” patients had significantly higher expression of the pathways “DNA Repair” and “HRD Through Homologous Recombination HRR”, supporting the hypothesis that this CLF describes a common DNA repair process predominant in CESC patients (**Fig. 2a**). Similarly, “*Factor 6 – T cell immune”* identified pathways, such as “Cytokine Signaling in Immune System”, and “PD-1 signaling”, that further strengthened the hypothesized function, while “*Factor 5 – B cell immune”* identified the pathways “Signaling by the B Cell Receptor (BCR)” and “Initial Triggering of Complement”. For primary-specific CLF “*Factor 12 – Head and Neck”*, the pathway “Keratinization and Formation of the Cornified Envelope” was highlighted. We further investigated significant pathways by considering the genes in these pathways that were outside the top 100 weighted genes for a CLF but still enriched in the respective high CLF patient subsets (**Fig. 2b**). For example, *PDCD1*, in the pathway “PD-1 signaling”, was specifically weighted in the corresponding “high” CLF population, in this case, “*Factor 6 – T cell immune”*. The differential expression of CLF-associated pathways, as well as the specificity of expression of pathway-associated genes in high-CLF patients, were validated in CPTAC HNSC and LUSC patients (**Supp. Fig. 6**). These results provide support that the CLFs capture biologically meaningful transcriptional programs.

To investigate whether distinct subsets of patients were associated with high weights for immune CLFs “*Factor 5 – B cell immune”* versus “*Factor 6 – T cell immune”*, we analyzed the expression of canonical immune cell-type markers in the mixed-primary populations of patients with relatively high weights for both CLFs, either CLF alone, or neither (**Supp. Fig. 7a**). Markers for B cell phenotypes, including Bregs and plasmablasts, as well as antibody-dependent cellular cytotoxicity (ADCC)^32^, and T cell phenotypes tended to be most highly expressed in patients high for both “*Factor 5 – B cell immune”* and “*Factor 6 – T cell immune”* (**Supp. Fig. 7b**). Patients high for only one CLF, either “*Factor 5 – B cell immune” or “Factor 6 – T cell immune”* were distinguished by their expression of T and B cell markers. High “*Factor 5 – B cell immune”*, low *“Factor 6 – T cell immune”* patients had greater expression of *CD1D*, a Breg marker, *CD27*, a memory B cell marker, as well as *IGHD* and *IGHM*, naïve B cell markers. Low “*Factor 5 – B cell immune”* and high “*Factor 6 – T cell immune*” patients were higher in expression of *CD86* and *CD80*, co-stimulatory markers, ADCC markers like *CCL3, CCL4*, and *CCL4A2*, as well as antigen presentation genes, like *CD68, HLA-DOA, HLA-DRB1*, and *CD163*. Furthermore, the hypothesized processes captured by these CLFs were supported by the immune cell-type compositions of each sample inferred by CIBERSORT^33^, with B cells significantly enriched in samples high in “*Factor 5 – B cell immune”* (**Supp. Fig 7c,d**) and T cells significantly enriched in high “*Factor 6 – T cell immune”* samples (**Supp. Fig. 7c,e**). Additionally, myeloid cells, such as macrophages and mast cells, were more frequent in patients low in both immune programs (**Supp. Fig. 7c,f**). Thus, consistent with other studies, the latent factor analysis highlights distinct, functional B and T cell responses as key, across-primary features of anti-tumor immunity.

### Distinct transcriptional CLFs are associated with clinical biomarkers and cell type gene signatures

Our observation that CLFs align with salient biological programs does not indicate whether these programs are useful for clinical decision-making. To further investigate the clinical meaning of the CLFs, we assessed the correlation of each CLF with known clinical variables, such as HPV status, smoking history, oncogenic mutation presence, and gene expression signatures (**Fig. 3a**). “*Factor 2 – HPV Positive”* and “*Factor 6 – T cell immune”* are enriched for HPV positivity in both HNSC and CESC. “*Factor 6 - T cell immune”* is enriched for patients with any CASP8 mutation. Additionally, this result is reproduced in HNSC CPTAC patients with CASP8 mutations. Interestingly, “*Factor 9* – *Smoking status”* is enriched for current smokers, whereas “*Factor 6 - T cell immune”* is negatively correlated with smoking history in HNSC. The CLF “*Factor 6 - T cell immune”* is also highly correlated with a continuous gene signature variable representing the ratio of cytotoxic interferon gamma to immunosuppression^17^. We also found that “*Factor 11 – Collagen Remodeling”* was highly correlated with a gene signature associated with localized invasion^34^, whereas “*Factor 1 – DNA Repair/Cervical”* was inversely correlated with it, suggesting less localized invasion in tumors with high CLF *1* weights. In the set of samples restricted to stage 4 cancers, which controls for the independent effect of staging on overall survival, “*Factor 11 – Collagen Remodeling”* was negatively correlated and with overall survival in both lung and head and neck cancers, whereas “*Factor 6 - T cell immune”* was positively correlated with overall survival in lung and cervical cancers.

**Figure 3:**
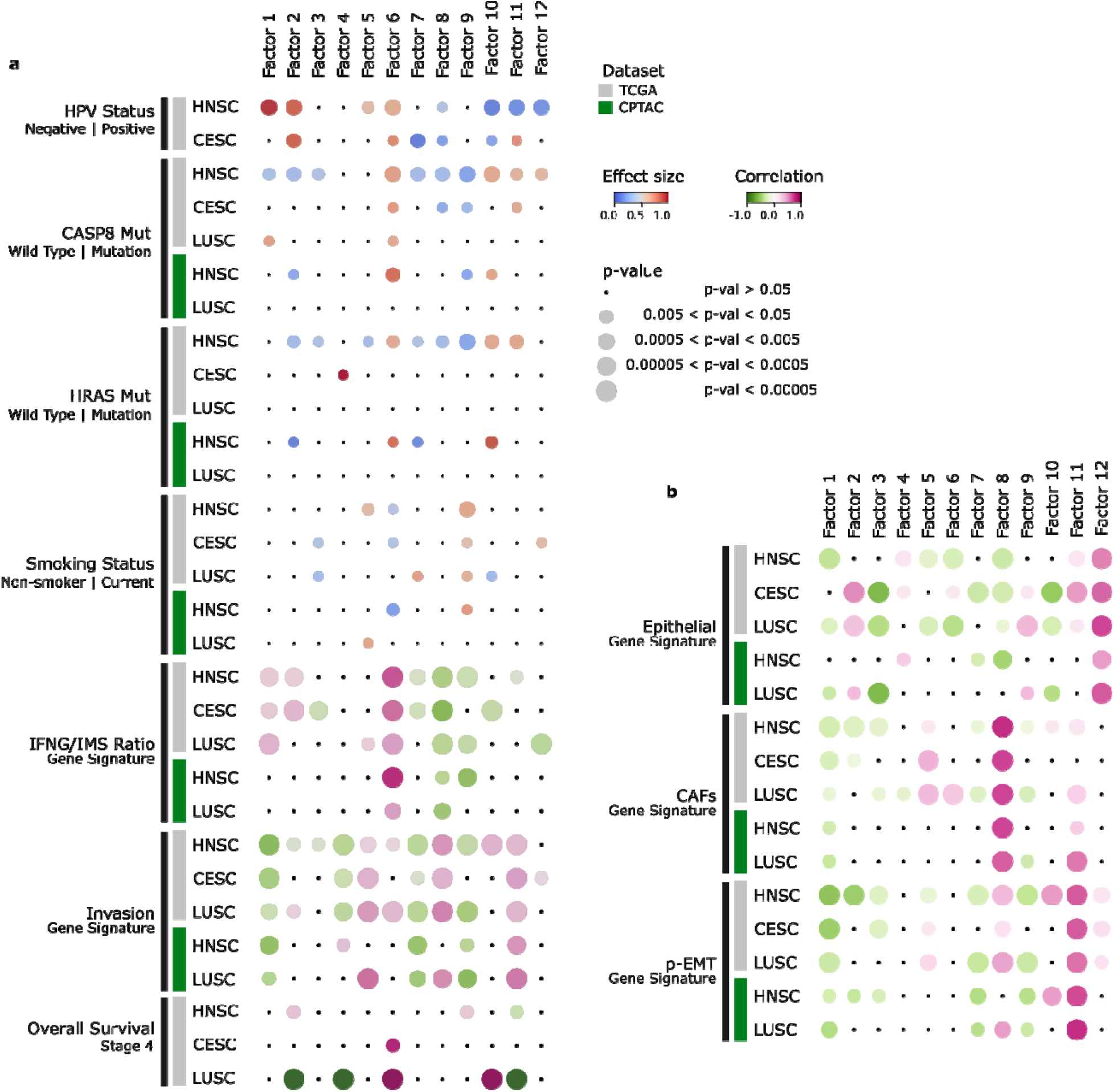
Distinct latent factors align with clinical biomarkers and tumor cell type gene signatures. **a**, Dot lot shows the association of several biomarkers (labels on left) with latent factor weights (columns), separately for each primary type and dataset (rows). Associations for binary biomarkers (HPV status, CASP8 gene mutation, HRAS gene mutation, and history of smoking) are indicated by the Mann-Whitney U statistic (color); effect size >0.5, in red (versus effect size <0.5, in blue), indicates higher (versus lower) than expected CLF values in patients with the biomarker. Associations for continuous biomarkers (IFNG/IMS gene signature ratio, invasion gene signature, and overall survival at stage 4) are indicated by Spearman’s rho correlation (color; pink, positive; green, negative). Dot size indicates p-value. **b**, Dot plot shows associations of cell type gene signatures (labels on left) with latent factor weights (columns), sepa ately for each primary type and data set (rows), by Spearman’s rho correlation (color); dot size indicates p-value.

Previous single-cell RNA-seq analysis of HNSCs identified gene markers for different cell types, including epithelial cells and fibroblasts, as well as markers of partial epithelial-to-mesenchymal transitions (p-EMT), associated with metastasis^35^. We created gene signatures for each set of cell-type specific genes and scored each sample using a previously described approach^17^. The epithelial gene signature scores were significantly correlated with “*Factor 12 – Head and Neck”* weights, suggesting that this CLF may represent a baseline HNSC signature, while “*Factor 8 – Fibrosis”* weights were significantly correlated with the fibrosis signature (**Fig. 3b**), underscoring our interpretation of that CLF. Additionally, the p-EMT scores were significantly correlated with “*Factor 11 – Collagen Remodeling”* weights, further supporting the hypothesis that this CLF is related to tissue remodeling and a propensity for invasion. Taken together, these results suggest that the CLFs capture several established features of SCC biology and disease biomarkers and demonstrate their significant heterogeneity and associations across patient populations and disease outcomes.

### Transcriptional CLF weights can be predicted from tumor histology

Recent studies have shown that machine learning methods can use tumor histology to predict key disease characteristics, including grade^36–38^, immune infiltration^39–41^, HPV status^42^, mutation status^19,43–45^, expression of individual genes^20,46,47^, and established multi-omic features^41^, but have not attempted to predict *de novo* learned, complex transcriptional features, such as our CLFs. To assess whether and how each CLF was histologically encoded, we trained a deep learning network to predict binarized CLF status (high or low) from tumor histology images (**Fig. 4a**). As before, the binarized high or low CLF label reflected whether the sample had a weight (amplitude) above the 75^th^ percentile or below the 60^th^ percentile, respectively, for that CLF (**Fig. 4b,c**). Prediction accuracy of the model was then assessed by the area under the curve (AUC) of the receiver operating curve (ROC). Models were trained separately on data from each primary type for which sufficient (50 or more) examples from each class (high or low) were present.

**Figure 4:**
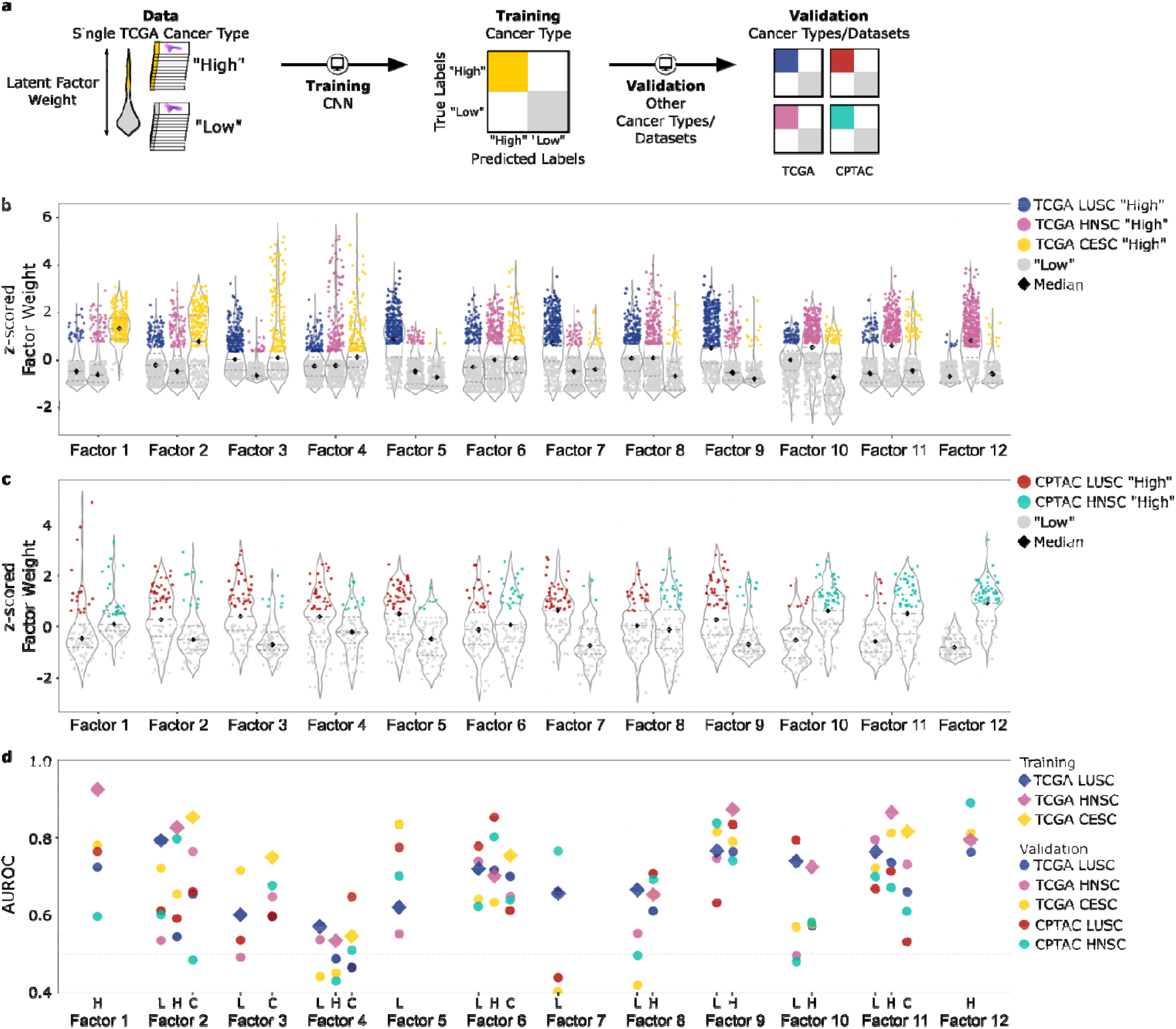
Neural networks were trained to successfully detect transcriptional latent factors in tumor histology. **a**, Schema for training and validation of deep learning models that predict high/low status of CLFs directly from tumor histology images. **b**, The TCGA data used to train the models are presented in violin plots showing the distributions across patients of the z-scored factor weights (y axis), z-score performed across patients per CLF in each dataset separately, separated visually by latent factor and primary cancer type (x axis). Data points are colored by bin, i.e., the high/low status used for training: patients with “high” (>75^th^ percentile) factor weights are colored according to their primary cancer type; those with “low” (<60^th^ percentile) weights are shown in light grey; percentiles are calculated per CLF across all TCGA patients and primary types. Internal boxplot shows median (thick dashed line), and first and third quartiles (thinner dashed lines). **c**, CPTAC validation data is presented in violin plots, analogous to those shown for TCGA (in b). **d**, Results of latent factor status predictions from tumor histology. For each latent factor and primary type with sufficient training data, a separate model was trained (x axis labels; L: lung; H: head and neck; C: cervical). Training results (diamonds, colored by training primary type) represent the average of the area under the receiver operating characteristic curve (AUROC) (y axis), where the curves are based on varying the threshold for tile-level calls, of 3 *k*-fold cross-validation rounds for a model with optimized hyperparameter settings. Trained models were validated on the other primary types and datasets; validation results (circles, colored by validation primary type and dataset) represent the AUROC of the model performance.

The results were strong, with prediction AUCs of at least 0.7 in TCGA for at least one primary type for 7 CLFs (i.e., “*Factor 1 – DNA Repair/Cervical”, “Factor 2 – HPV Positive”*, “*Factor 3 – Epithelial cells”, “Factor 6 – T cell immune”*, “*Factor 9 – Smoking status”*, “*Factor 10 – Metabolism”*, “*Factor 11 – Collagen Remodeling”*, and “*Factor 12 – Head and Neck”*) (**Fig. 4d**). For instance, “*Factor 1 – DNA Repair/Cervical”* was trained in TCGA HNSC with an average AUC (aAUC) of 0.92 across training and validation subsets. (It could not be trained on TCGA CESC due to insufficient “low” samples.) The model subsequently validated in other primary tissues (TCGA LUSC AUC: 0.72; CESC AUC: 0.78) and datasets (CPTAC LUSC AUC: 0.76; HNSC AUC: 0.60).

“*Factor 6 - T cell immune”* status was also accurately predicted, with training aAUCs of 0.70-0.75, and validation AUCs of 0.61-0.85. A few CLFs were not predicted as accurately (“*Factor 5 – B cell immune”*, “*Factor 7 – Lung”*, and “*Factor 8 – Fibrosis”*, with training aAUCs of 0.6–0.7). The specific, low accuracy (aAUCs of 0.55–0.57) for “*Factor 4 – Cellular stress”* may reflect its intracellular program.

While three performant models for “*Factor 11 – Collagen Remodeling”* were trained, one in each primary cancer type (TCGA LUSC aAUC: 0.76; HNSC aAUC: 0.86; CESC aAUC: 0.82), each model translated with a different degree of success to other primary types and datasets (**Fig. 4d**). The LUSC trained model performed similarly well in validation in other tissues (TCGA HNSC AUC: 0.79; CESC AUC: 0.72) and datasets (CPTAC HNSC AUC 0.70; LUSC AUC: 0.67), suggesting that the visual features learned by training on TCGA lung data generalize to cervical and to head and neck primary types, as well as to CPTAC. In contrast, while the CESC trained model had a similar aAUC, its performance dropped in validation (TCGA HNSC AUC: 0.73; LUSC AUC: 0.66), particularly in the CPTAC dataset (CPTAC HNSC AUC: 0.61, LUSC AUC: 0.53), suggesting that this model learned cervical- and TCGA-specific image features. Collectively, these results demonstrate that our deep learning models were trained to accurately detect high or low factor weights for many CLFs, directly from tumor histology.

### Synthetic histological images reflect CLF phenotypes

While predictive of CLF scores, the deep learning predictions of gene expression scores do not indicate which features are responsible for that prediction to facilitate human interpretation. To explore the image features that support the model-based CLF classification of tumor histology, we used a conditional generative adversarial network (cGAN) to generate synthetic H&E stained histologic images prototypical of a given class, as well as images that smoothly transition between classes^48^ (**Fig. 5**). Specifically, for each primary cancer type and a given CLF, we used the cGAN to generate pairs of images reflective of the CLF class labels in that cancer type, one version in a “high” CLF style and another in a “low” CLF style (**Fig. 5a)**. Then we interpolated between the two images to infer intermediate histologic states, reflecting the continuous nature of the original CLF embedding. This process allowed a pathologist, blinded to the hypothesized biological function of the CLF, to then annotate histologic features perceived as differential between the “high” and “low” CLF style synthetic images, thereby identifying interpretable image features potentially responsible for the “high” versus “low” CLF model predictions (**Supp. Table 2**).

**Figure 5:**
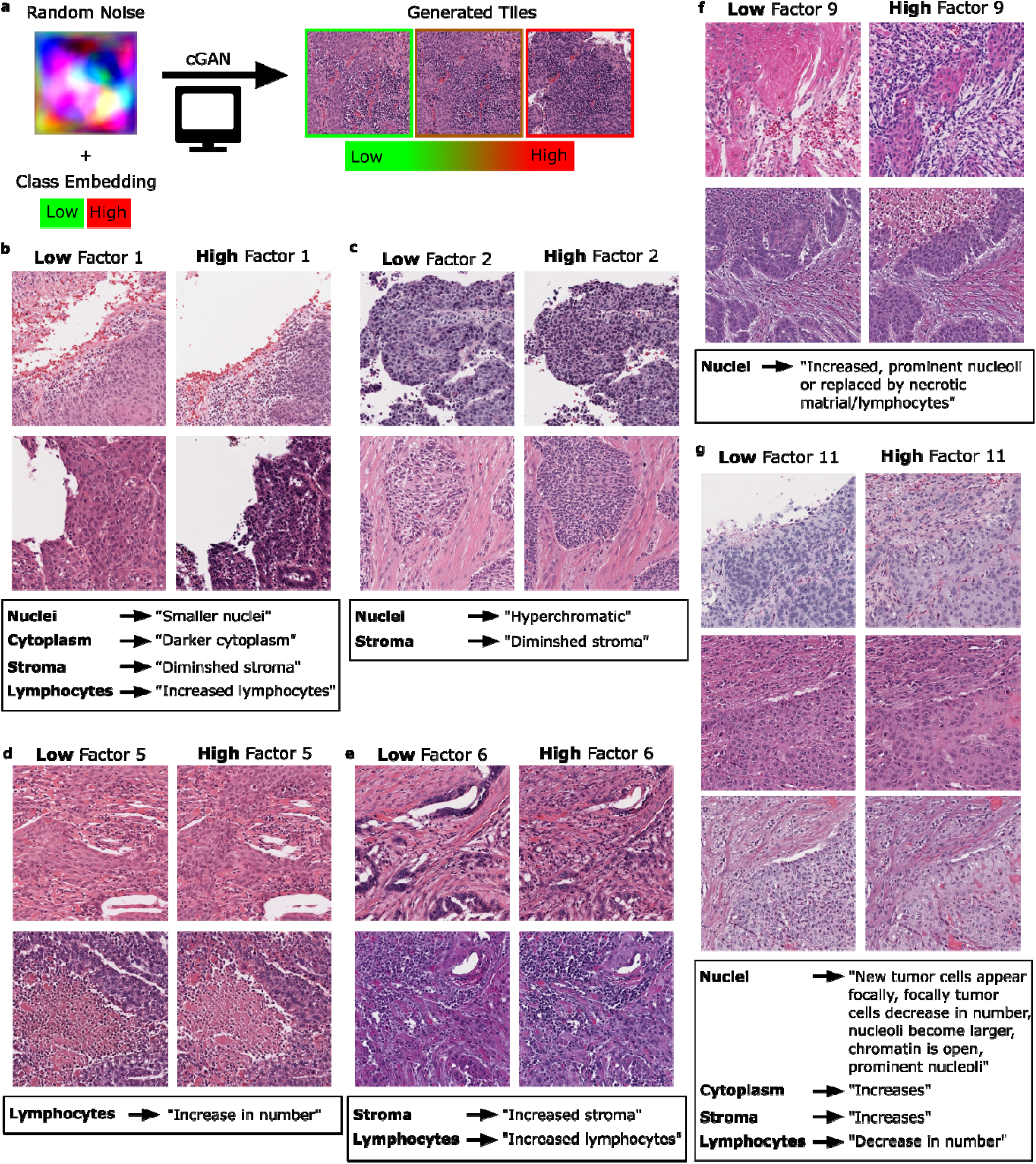
Generated synthetic image tiles illustrate histologic features associated with transcriptional latent factor status. **a**, Schema for generating histology image tiles corresponding to the high/low status of a latent factor, using a conditional generative adversarial network (cGAN). **b–g**, For latent factors 1, 2, 5, 6, 9, and 11, respectively, tiles illustrate distinct histologic features, with pathologist characterizations (bottom annotations), associated with low (left column) versus high (right column) factor status, for two random seeds (rows).

Of the most performant models, “*Factor 6 - T cell immune”* and “*Factor 1 – DNA Repair/Cervical”* had especially notable differential histologic features. The pathologist reported that synthetic images high in “*Factor 1 – DNA Repair/Cervical”* showed smaller nuclei, darker cytoplasm, diminished stroma, and increased lymphocytes (**Fig. 5b**). Images high in the CLF “*Factor 2 – HPV Positive”* similarly showed diminished stroma, but with hyperchromatic nuclei (**Fig. 5c**). Notably, while these CLFs are high in similar patient populations, predominantly CESC and HNSC, the histologic features reflecting CLF presence are distinct. Similarly, images high in “*Factor 5 – B cell immune”* showed increased lymphocytes (**Fig. 5d**), as did those high in “*Factor 6 – T cell immune”* (**Fig. 5e**), but the former additionally exhibited increased stroma. Unlike the CLFs with established histologic correlates, images high in “*Factor 9* – *Smoking status”* were annotated with apparently novel nuclear and cell-type traits (**Fig. 5f**). Images high in “*Factor 11 – Collagen Remodeling”* (**Fig. 5g**) showed new tumor cells appearing, as well as increased cytoplasm, increased stroma, and a decrease in the number of lymphocytes. These data, comprising both standard and novel histologic features, provide potential explanations for the image features each deep learning model learns to associate with a CLF. Additionally, the histologic features align well with the hypothesized biological programs associated with the CLF, supporting the notion that a common latent biological process underlies both the genomic and histologic data features.

## Discussion

Our study demonstrates a role for unsupervised latent factor analysis of gene expression to uncover key, histologically encoded features of SCC biology. We provide a systematic analysis of the results of a Bayesian non-negative matrix factorization, demonstrating that transcriptional CLFs capture not only primary-specific, but also cross-primary, programs associated with disease etiology, as well as known genomic and clinical features of the disease. For example, CLFs “*Factor 5 – B cell immune”* and “*Factor 6 – T cell immune”* highlight the existence of diverse immune phenotypes within SCCs. Consistent with prior work showing that T-cell infiltration of HNSC tumors is associated with favorable outcomes^49^, “*Factor 6 – T cell immune”* weights are correlated with increased overall survival in stage 4 SCCs. From this unsupervised factorization we identified CLFs that capture the significant heterogeneity in transcriptional programs related to specific, individual factors like disease etiology, cell composition, propensity for invasion, and primary organ type among SCCs.

In addition to profiling the genomic and clinical correlates of CLFs, we investigated the extent to which gene-expression–derived CLFs could be detected directly in tumor histology, thereby increasing their clinical translation potential. we found that CLFs representing a variety of biological processes were robustly detected in tumor histology, results that we validated both across different SCC primary tissues and datasets. The most performant models included those detecting “*Factor 1 – DNA Repair/Cervical”, “Factor 2 – HPV Positive”*, “*Factor 3 – Epithelial cells”, “Factor 6 - T cell immune”*, “*Factor 9 – Smoking status”*, “*Factor 10 – Metabolism”*, “*Factor 11 – Collagen Remodeling”*, and “*Factor 12 – Head and Neck”*. While previous studies predicted other types of genomic features from digital pathology, our study provides the first evidence that *de novo* inferred transcriptional programs can be detected robustly in tumor histology, and that the histological features that predict such programs may generalize across primary tissues and datasets.

To further investigate the histologic features corresponding to CLF detection, we utilized cGANs to generate example images representing “low” versus “high” CLF classes and to interpolate between the classes. Histologic features of synthetic images were termed by a pathologist blinded to the hypothesized biological function of the CLFs. These annotations revealed coherent phenotypes, which also corresponded to the hypothesized biological processes, for “*Factor 1 – DNA Repair/Cervical”*, “*Factor 2 – HPV Positive”, “Factor 5 – B cell immune”*, “*Factor 6 – T cell immune”*, “*Factor 9* – *Smoking status”*, and *“Factor 11 – Collagen Remodeling”*. Hence, we demonstrate a new approach that takes advantage of computer-generated histologic images to assess histologic features used by deep learning models for classification. The coherent, expected histologic features that were identified by pathologist review support the assertion that the histology and genomics features jointly reflect relevant, underlying SCC biology. This use of genomic technologies to inform histology-based patient classification also demonstrates a potential approach for translating complex gene expression-derived features into interpretable histologic features.

The particular set of CLFs we identified represents one of many possible nonnegative factorizations of the same gene expression matrix. Moreover, latent factorization methods are not designed to detect rare biological programs, which may be treated similarly to technical noise. To evaluate the sensitivity of our CLFs to analysis and model parameter choices, we used multiple data normalization schemes, factorization approaches, and model parameters, including different numbers of factors, to compute other factorizations and compare results. The majority of the CLFs we describe above, including “*Factor 11 – Collagen Remodeling”* and “*Factor 5 – B cell immune”*, were detected robustly across other factorizations; however, some partially overlapping CLFs, such as “*Factor 6 – T cell immune”* and “*Factor 5 – B cell immune”*, were frequently combined into a single factor. The robustness of our results across parameter combinations, the correlation of CLFs with clinical and mutation data, and the detection of CLFs in tumor histology all suggest the factorization we used is a valid, appropriate solution that reflects meaningful variation in both the TCGA and CPTAC datasets.

Certain CLFs were not well detected in tumor histology, as evidenced by low AUC values. The challenge in their detection may be at least partly related to their biological functions; for instance, the intracellular program of “*Factor 4 – Cellular stress”* may simply not exhibit visual changes at the level of the tissue architecture captured by H&E slides. Additionally, to ensure that we identified predictive histologic features that generalized across SCC primary tissue types, we trained models separately within each primary, and did not train on a primary if there were fewer than 50 primary samples identified as either “high” or “low” for a specific CLF. This requirement excluded certain CLF and primary combinations from training, such as “*Factor 1 – DNA Repair/Cervical”*, which is “high” in most CESC samples. This restricted some potentially information-rich sets of slides to be used solely for validation.

Altogether, our results identify biologically salient, gene-expression–derived features of SCCs that are reflected in histology, as well as demonstrate a novel approach for translating these de novo transcriptomic features into interpretable histologic features. Beyond deciphering the latent biology of SCCs, histology-encoded CLFs may represent programs that are informative of treatment response and, in turn, shed light on the complex biological processes involved in therapeutic success.

## Methods

### Analysis Overview

All data were analyzed using publicly available software packages, in combination with custom code to analyze and plot results. Custom code will be publicly available at a Zenodo repository (upon publication).

### RNA-seq Data Acquisition and Processing

After removal of vendor-failed reads, RNA-seq raw reads were aligned to human GRCh38 reference genome using STAR (ver. 2.7)^50^ with the basic 2-pass mapping method. Gene quantification was performed using the STAR GeneCounts quantification mode with the GENCODE gene set (ver. 36).

TCGA and CPTAC patient samples of non-squamous cell origin were removed from TCGA-CESC, leaving only squamous cell carcinomas. The TCGA dataset was then TMM-normalized using edgeR (ver. 3.38.1)^51^, and CPTAC was subsequently filtered according to the TCGA-derived gene filtering list and TMM-normalized.

### Genomic Mutation Data

Genes considered for mutation status comparisons were selected from the OncoKB^52^ v4.0 database by focusing on “actionable mutations” in SCC. Only mutations with significant CLF associations were shown. Data was pulled using the cBioPortal API for R. For each gene, values were binarized into “mutation” and “wild type”, such that any mutation in the given gene was assigned to “mutation”, and no mutation, i.e., a match to the reference, was assigned to wild type.

### Bayesian Non-negative Matrix Factorizations of RNA-seq Data

Bayesian non-negative matrix factorization (NMF) was performed using CoGAPS (ver. 3.16.0)^26^ on log-scaled TMM-normalized TCGA RNA-seq data (specifically, log_10_(*x*+1) for TMM-normalized expression value *x*). The factorization presented and analyzed was performed using 12 CLFs, 1000 iterations (“nIterations”), and a random seed of 34. The function featureLoadings was used to exclude any genes not present in the CPTAC validation dataset, and the function projectR (ver. 1.12.0)^53^ was used to factorize CPTAC by projecting it into TCGA-derived latent space, i.e., to hold the feature loadings for each factor constant and compute factors weights (“sampleFactors”) for CPTAC samples.

CLFs were reordered for better visualization as a heatmap, using the slanted_orders function from slanter (ver. 0.2-0) to reorder the columns of the matrix returned by the sampleFactors function of CoGAPS.

To compare CLF weights across datasets, each CLF was z-scored across all primaries within a single dataset (either TCGA or CPTAC). Multi-dataset UMAP was computed from these values with default parameters. For a given CLF, a patient was classified as “high” or “low if the CLF sample weight was above the 75^th^ percentile or below the 60^th^ percentile, respectively, where percentiles were computed for each CLF across all primaries in a single dataset.

### GTEx Bulk Tissue Expression

The Genotype-Tissue Expression (GTEx) Project was supported by the Common Fund of the Office of the Director of the National Institutes of Health, and by NCI, NHGRI, NHLBI, NIDA, NIMH, and NINDS. The data used for the analyses described in this manuscript were obtained from the GTEx Portal on 03/13/2023.

### Pathway Analysis

The top 100 weighted genes from each CLF were fed unordered into the g:Profiler program g:GOSt (ver. E106_eg53_p16_65fcd97), where the statistical domain scope was the TCGA filtered gene list with default multiple test correction, g:SCS significance threshold. Results were then filtered to only include Reactome pathways with an adjusted p-value <0.01. Full gene lists for the pathways identified by g:GOSt were pulled from the Molecular Signatures Database (MsigDB)^54,55^ v7.5.1.

Average log-scaled gene expression was computed by taking the mean across all genes of log(x+1), where *x* represents for normalized TMM expression of a gene. For specific pathways, the average was computed over all genes listed in the Reactome pathway that were also included in the counts matrix. In contrast, average z-scored gene expression was computed by z-scoring normalized expression of each gene individually and then averaging across patients in the respective group.

### Gene Signature Scores

Gene signature scores were derived from a list of genes using the arithmetic mean of the log2-transformed gene list, as was described for the previously published IFNG/IMS gene ratio^17^.

### Slideflow

For histology-based CLF detection, a patient sample was annotated using the same “high” or “low” CLF labels as previously described, i.e., if its factor weight was above the 75^th^ percentile or below the 60^th^ percentile, respectively, of weights for that CLF. Patient samples with weights between the 75^th^ and 60^th^ percentiles were censored to create a greater distinction between the groups.

We trained deep learning classification models using Slideflow to implement an Xception architecture, using ImageNet pretrained weights and two hidden layers of width 1024, with dropout (*p* = 0.1) after each hidden layer. In Slideflow (https://doi.org/10.5281/zenodo.5703792) models were trained using the Tensorflow backend on 299-pixel histopathological image, representing non-overlapping tiles from H&E-stained tumor slides at 10X effective magnification (302 microns). Histopathological images were stain normalized using a modified Reinhard^56^ method with the brightness standardization step removed for computational efficiency. Training image tiles were augmented with random flipping and cardinal rotation, JPEG compression (50% chance of compression with quality level between 50-100%), and gaussian blur (10% chance of blur with sigma between 0.5-2.0). Binary categorization models were trained with cross-entropy loss. Models were first trained within each individual TCGA primary dataset (LUSC, CESC, or HNSC) for each CLF with a minimum of 50 samples in each binary category. To avoid learning features specific to data-collection sites, we performed site-preserved cross-validation^57^, using the parameter k-fold-preserved-site, for training for hyperparameter selection. The learning rates and number of epochs were varied across training models, and hyperparameters were selected from the best performant model across either TCGA LUSC, HNSC, or CESC, where performance was evaluated by averaging across 3 k-folds of the area under the receiver operating characteristic curve (AUROC). The curves for AUROC were based on varying the threshold for tile-level calls. A final model was then trained across the full dataset, either TCGA LUSC, HNSC, or CESC, and tested on the other primary TCGA datasets and CPTAC datasets.

### cGAN training

Our cGAN architecture is an implementation of StyleGAN2^58^, minimally modified to interface with the histopathology deep learning package Slideflow and to allow for easier interpolation in the embedding space^48^. For a given CLF, a cGAN was conditioned on the category of CLF weight (“high” versus “low”), in combination with the category of primary cancer type (“LUSC” versus “HNSC” versus “CESC”).

The cGAN was trained on 2 A100 GPUs for 25,000 kimg (25 million total images), initialized with random weights, and was stopped at a particular time point due to model divergence with further training. All cGANs were trained with an R1 gamma of 1.6384, batch size of 32, and using all available augmentations from StyleGAN2.

### cGAN class and layer blending

To create class-conditional images, cGAN class labels were projected into an embedding space before conditioning the network using the projection learned during training. Thus, after training, each class label has a single associated embedding vector. To create class-blended images, we perform a linear interpolation between class embeddings and use these interpolated embeddings for network conditioning while holding the cGAN seed constant^48^. We create layer-blended images by passing different class embeddings to each cGAN network layer while holding the cGAN seed constant.

### Classifier concordance

To assess classifier concordance for a cGAN and an associated classifier, the cGAN was used to generate an image for each class (i.e., “high” and “low”) using the same seed. The generated images were center-cropped, resized, and color-normalized to match the same histologic magnification and normalization as the associated classifier, and the classifier created predictions for each image. Predictions were considered “strong” if the post-softmax value was greater than 0.75 for the predicted class, and “weak” if the post-softmax value for the predicted class was less than 0.25. A given seed was defined as “strongly concordant” if (1) the classifier predictions matched the cGAN class labels for both images, and (2) the predictions were both strong. Only strongly concordant synthetic images were given to a pathologist for characterization.

### Pathologist assessment of cGAN images

A pathology fellow reviewed a subset of strongly concordant, synthetic histologic images, as well as the class blended videos interpolating from the “low” to “high” CLF states in each primary, with sufficient examples to characterize changes to the nuclei, cytoplasm, stroma, and lymphocytes. The pathologist reviewed the images in a blinded fashion without knowledge of the associated cGAN labels. Lossless, PNG images were viewed at the full resolution. The pathologist described histologic differences between each image pairs and recurrent, consistent descriptions were pulled as summary features (**Supp. Table 2**).

### Statistical analysis

Statistical tests for group comparisons included the *t*-test (**Fig. 2a, Supp. Fig. 6a**), and Mann-Whitney U-test (Fig 3a, Supp Fig 7d-f). Correlations were performed using Spearman rho tests (**Fig. 1c, Fig. 3a,b**, **Supp. Fig. 3**), and the p-values shown were Bonferroni corrected for testing 12 CLFs (**Fig. 1c,d**, **Fig. 2a**).

---

**Supplementary Figure 1:**
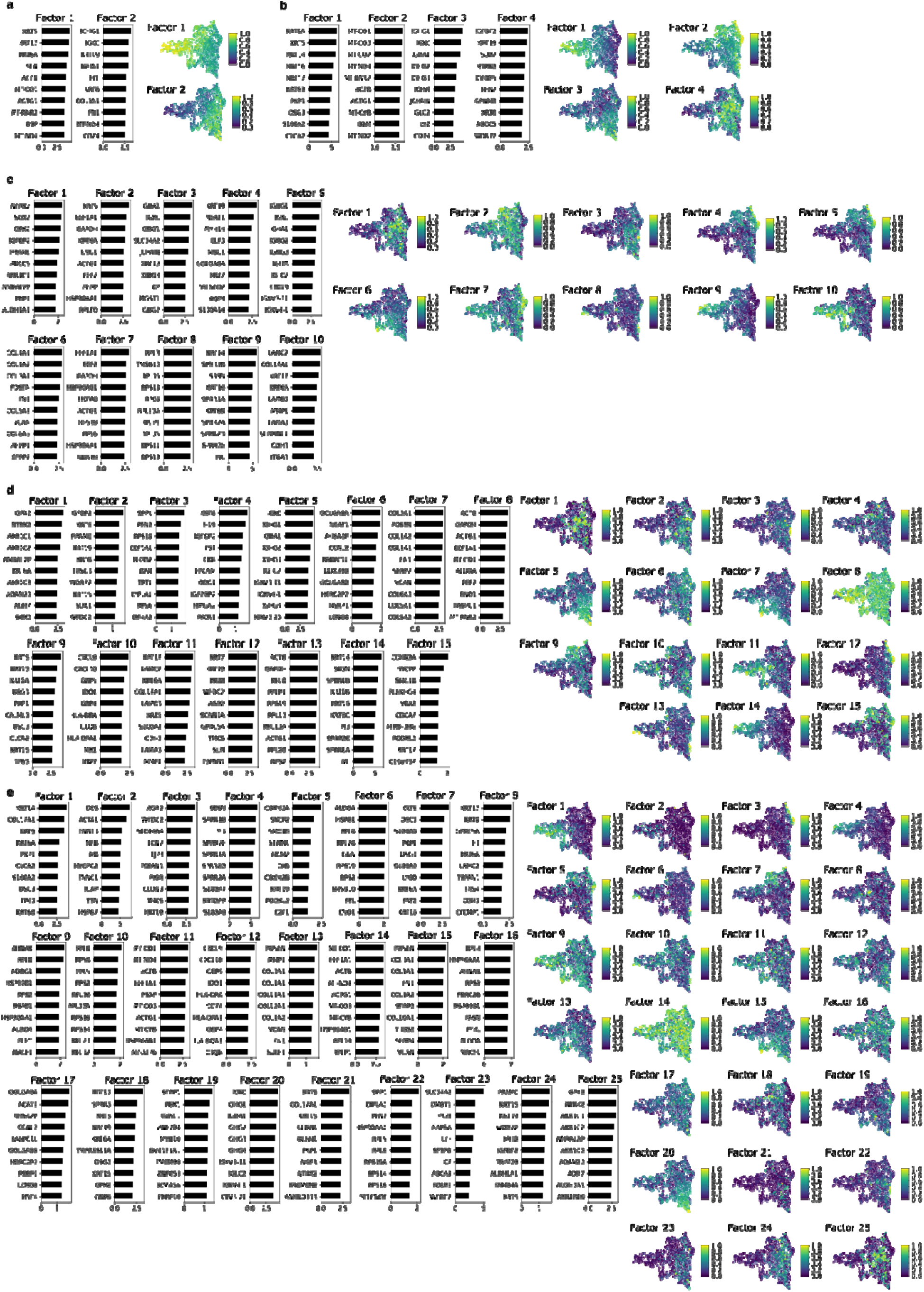
Selection of the number of CLFs. **a**, For a model with *k=*2 CLFs, for each factor, bar chart indicates top-weighted genes, and UMAP, created from the normalized TCGA transcriptomic data, shows patient factor weights (color). b–e, Analogous plots for models with varying numbers of CLFS: *k*=4 (**b**), *k*=10 (**c**), *k*=15 (**d**), and *k*=25 (**e**).

**Supplementary Figure 2:**
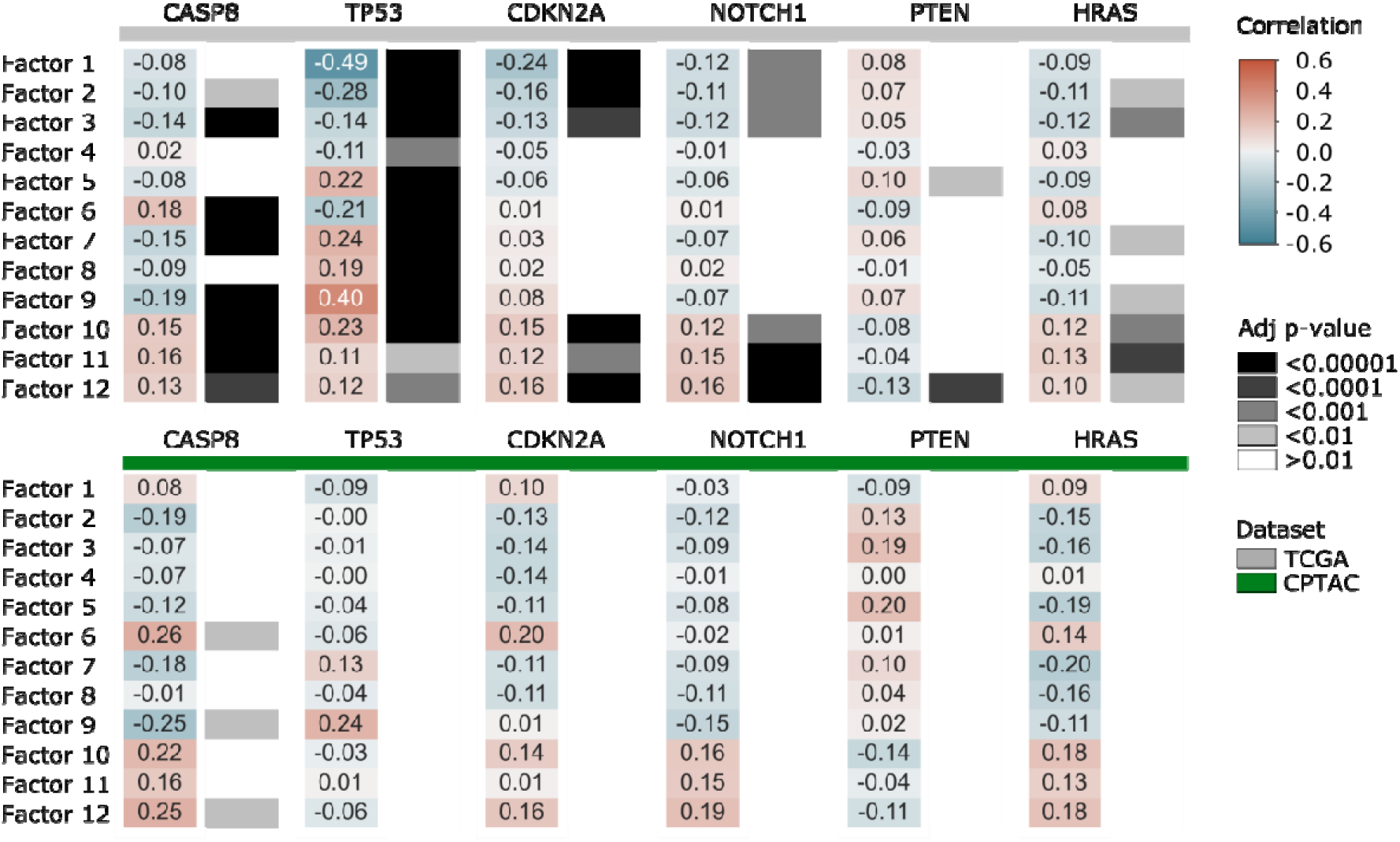
Transcriptional latent space factorization identifies mutation-specific variation. Plots show Spearman’s rho correlations (color), with Bonferroni-adjusted p-values (gray scale), between latent factors (rows) and mutation status for several genes (headings) in each dataset.

**Supplementary Figure 3:**
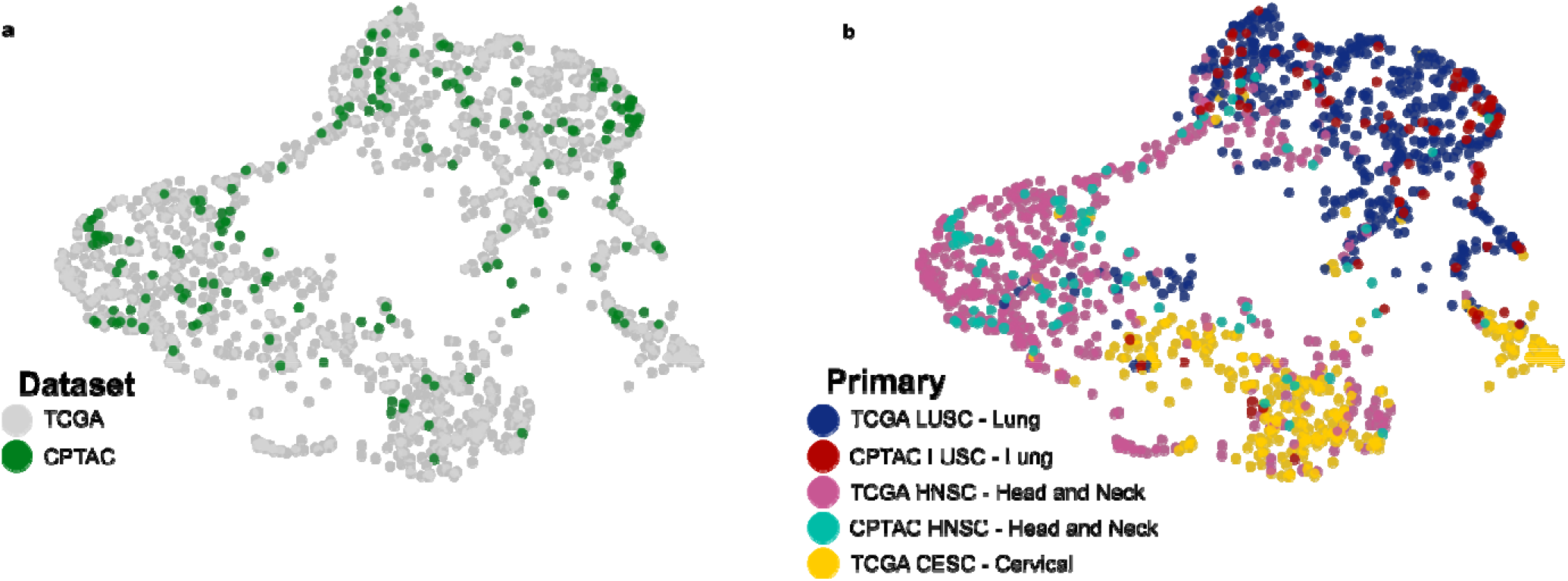
CPTAC data projection into the latent factor space inferred by CoGAPS from TCGA. a,b, A UMAP embedding was created from the CLF coordinates of TCGA patients, as inferred by CoGAPS, and of CPTAC patients, by projecting their transcriptomic data into the TCGA-defined CLF space. Patients (dots) are colored by dataset of origin (**a**), or by both primary cancer type and dataset (**b**).

**Supplementary Figure 4:**
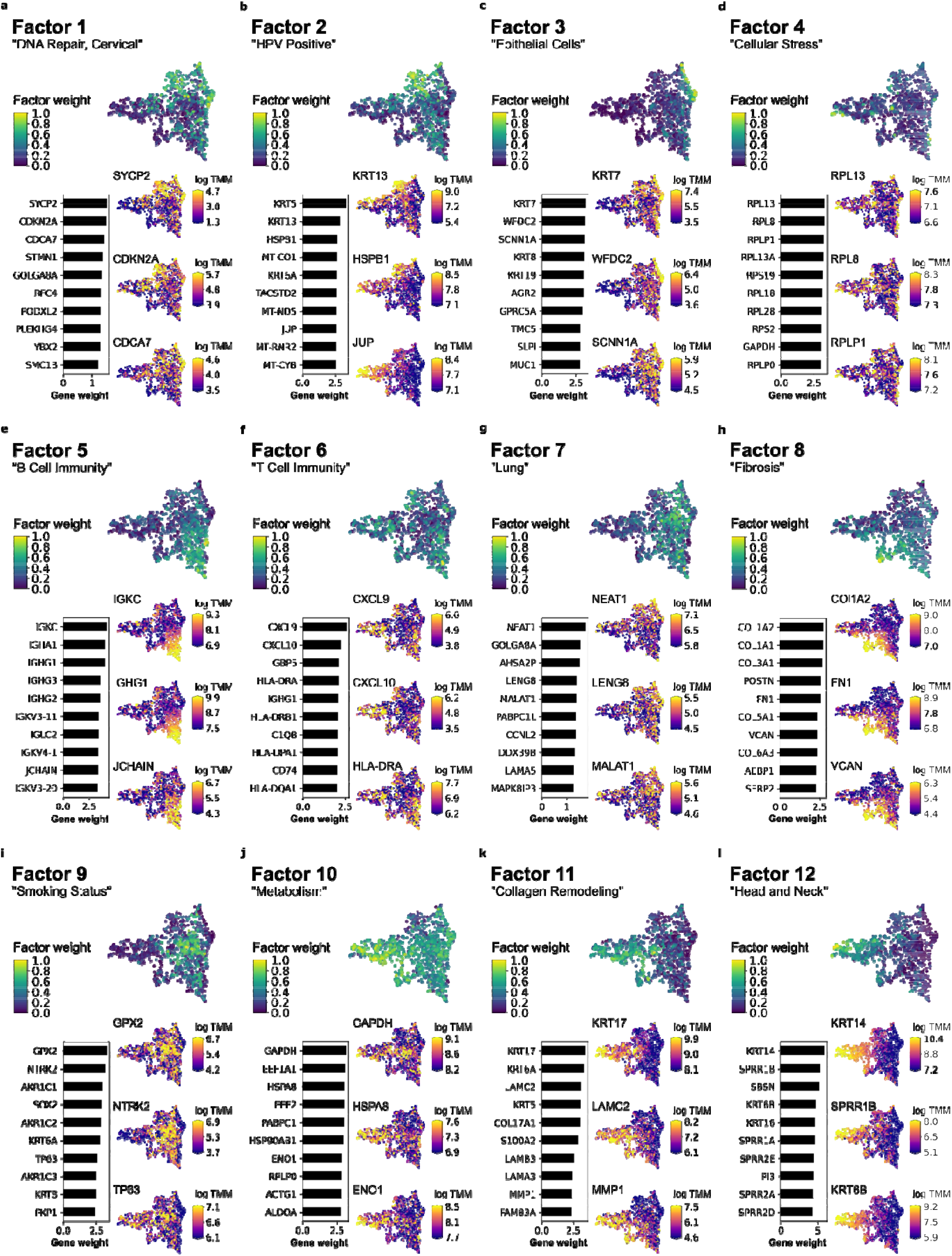
Latent factors are associated with salient features of SCC biology. **a**, For Factor 1, UMAPs (as in Supp. Fig 1) show patients (dots) colored by factor weight (top) or by expression (log TMM) of 3 highly weighted genes (right column); bar plot (left) shows top 10 genes (rows) by factor weight (x axis). **b–l**, Analogous plots are shown for Factors 2–12, respectively.

**Supplementary Figure 5:**
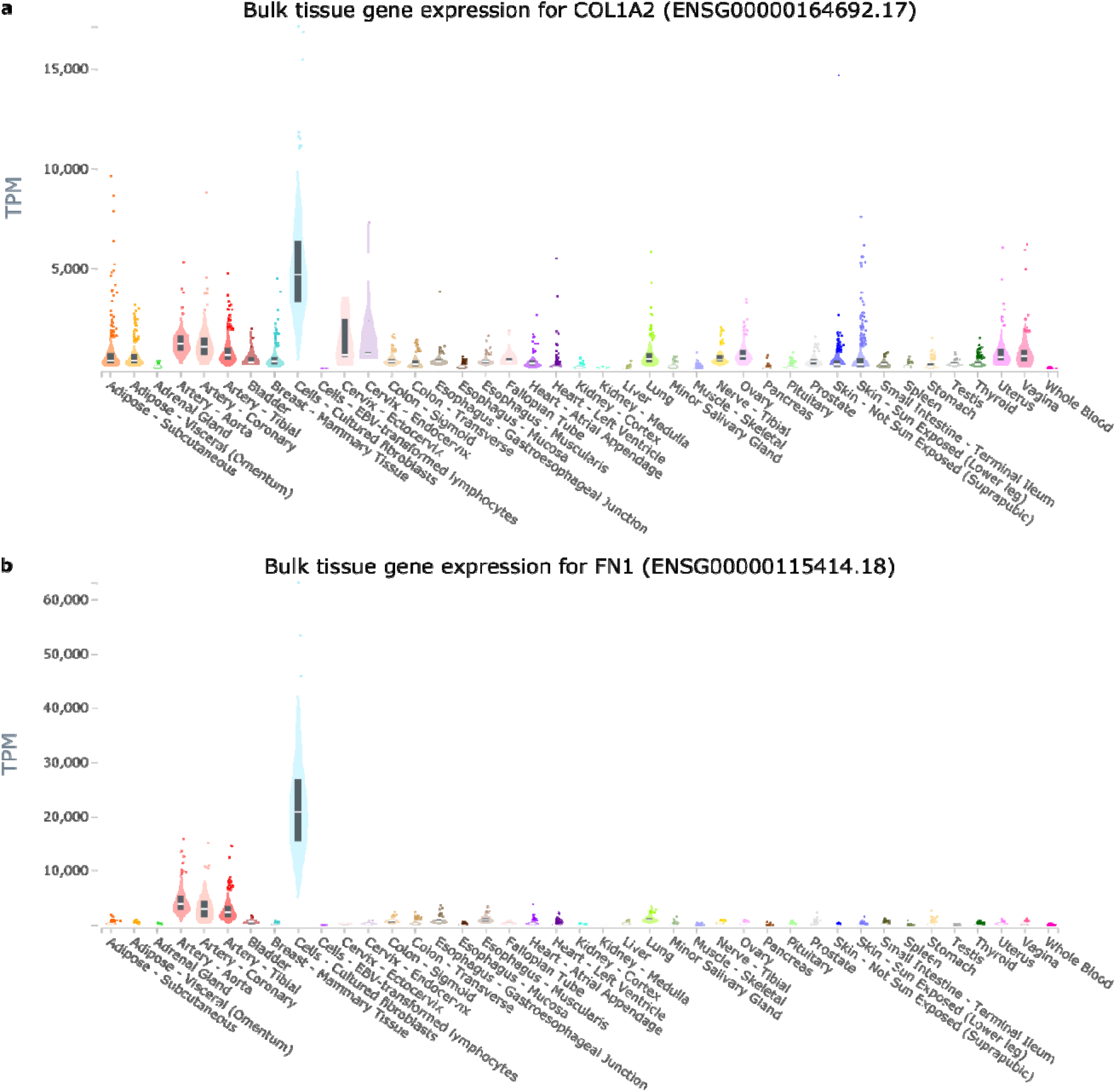
Factor 8 genes are highly expressed in fibroblasts. a,b, Violin plots show the distributions of expression values (TMM, y axis) in normal tissues (x axis) of the genes COL1A2 (**a**) and FN1 (**b**), both highly weighted in Factor 8, according to the genotype-tissue expression (GTEx) project. Internal boxplot shows the median (white line), and the first quartile and third quartile (bottom and top ends of thicker box).

**Supplementary Figure 6:**
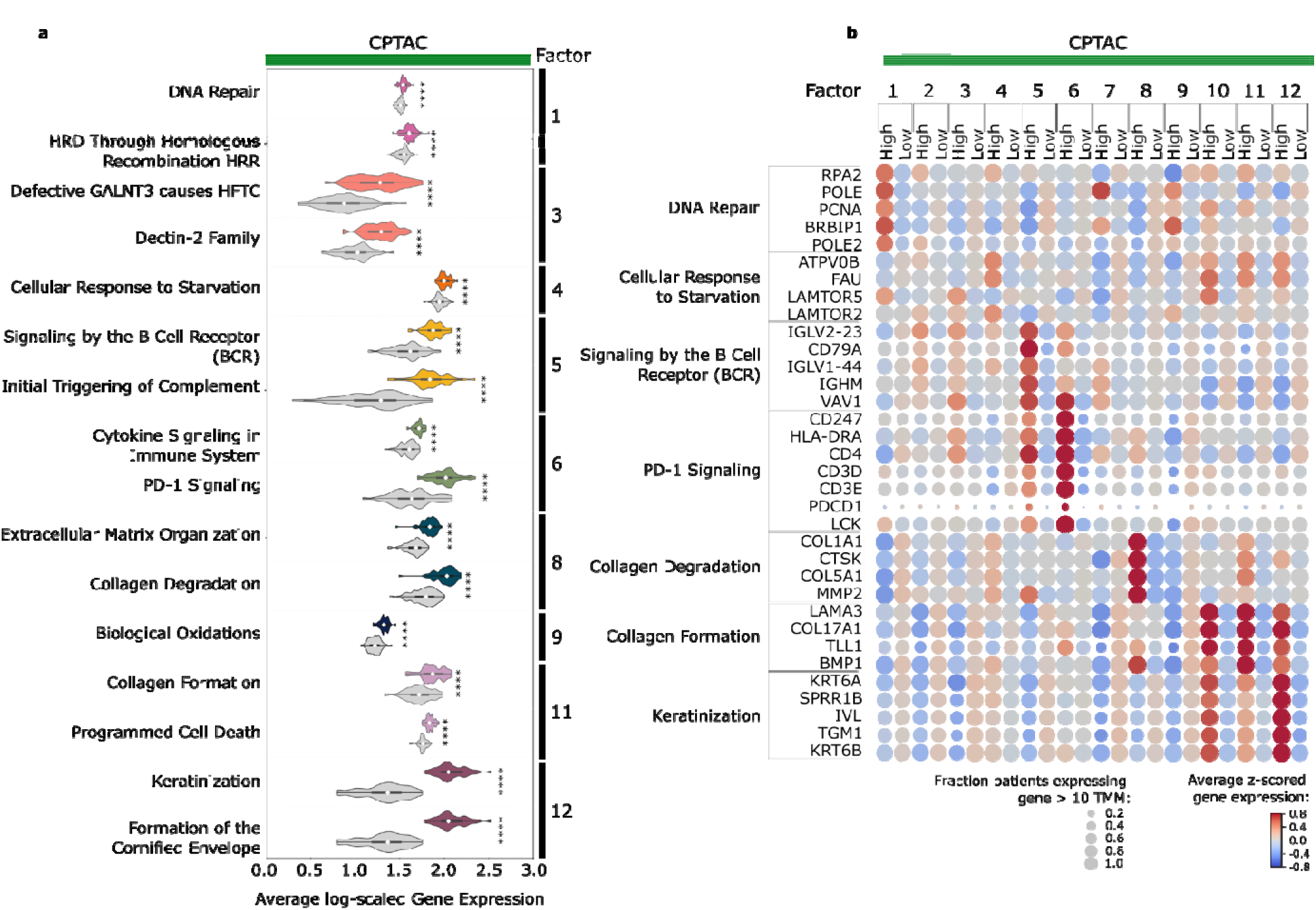
Latent factor genes are enriched in distinct, cancer-associated pathways. **a**, Violin plots show the distributions of the average log expression of all genes in a pathway (row label) that was identified as enriched in genes associated with the indicated latent factor (label on right), across CPTAC patients who have high (in color) or low (in gray) weights for that factor. Internal boxplot shows the median (white diamond), the first quartile and third quartile (left and right ends of thicker box), and the minimum and maximum (left and right ends of thin line). Distributions compared by *t*-test (**** *P* < 0.0001) **b**, Dot plot shows the average expression (color) of individual genes z-scored by expression across the dataset in CPTAC patients, grouped by high or low factor weight (columns, annotations above), for select genes (rows) in enriched pathways (annotations on left). Dot size indicates the fraction of patients in the group expressing the gene.

**Supplementary Figure 7:**
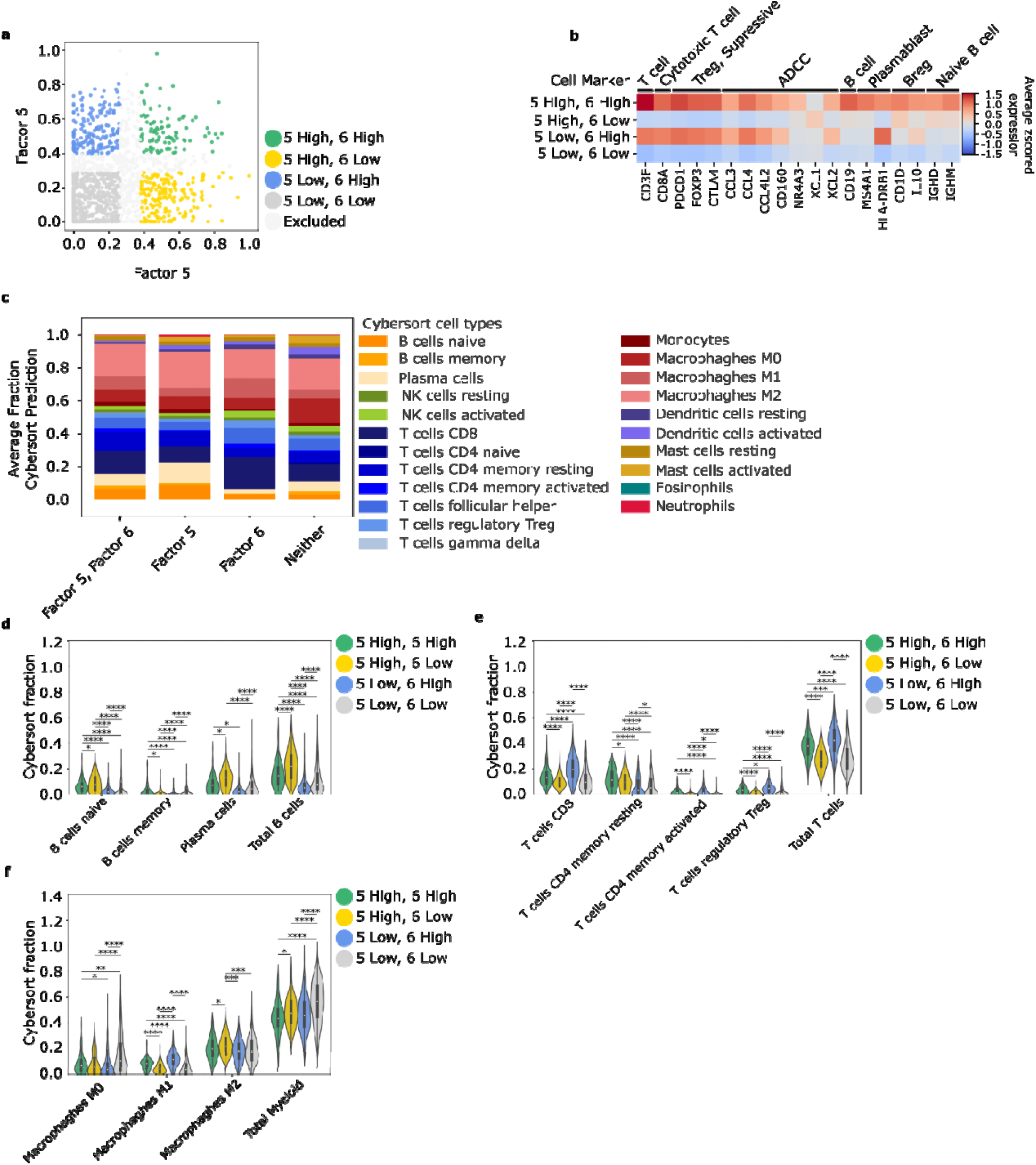
Factors 5 and 6 identify both distinct and common immune features of tumors. **a**, Scatterplot shows patients (dots) by their weights for “Factor 5 – B cell immune” (x axis) and “Factor 6 – T cell imune” (y axis), colored by their status as “high” or “low” for one, both, or neither factor. **b**, Heatmap shows average z-scored gene expression (color), individual genes z-scored within the TCGA dataset, of immune-cell marker genes (columns) in patient subsets (as in a). **c**, Bar charts show average fractions (y axis) of immune cell populations predicted by CIBERSORT, in patient subsets (as in a). d–e, Violin plots show distributions in patient subsets (as in a) of CIBERSORT-predicted fractions of B cell and plasma (**d**), T cell (**e**), and myeloid cell (**f**) types, with statistically significant differences indicated (Mann-Whitney U test, * *P* <0.05, ** *P* <0.001, *** *P* <0.0001, **** *P* <0.00001). Internal boxplot shows the median (white diamond), the first quartile and third quartile (bottom and top ends of thicker box), and the minimum and maximum (bottom and top ends of thin line).

## Notes

### Competing Interest Statement

The authors have declared no competing interest.

